# Formation of aggregates, icosahedral structures and percolation clusters of fullerenes in lipids bilayers: The key role of lipid saturation

**DOI:** 10.1101/2020.02.12.946152

**Authors:** Nililla Nisoh, Viwan Jarerattanachat, Mikko Karttunen, Jirasak Wong-ekkabut

**Affiliations:** Department of Physics, Faculty of Science, Kasetsart University, Bangkok 10900, Thailand; Computational Biomodelling Laboratory for Agricultural Science and Technology (CBLAST), Faculty of Science, Kasetsart University, Bangkok 10900, Thailand; Thailand Center of Excellence in Physics (ThEP Center), Ministry of Higher Education, Science, Research and Innovation, Bangkok 10400, Thailand; Specialized Center of Rubber and Polymer Materials for Agriculture and Industry (RPM), Faculty of Science, Kasetsart University, Bangkok 10900, Thailand; NSTDA supercomputer Center (ThaiSC), National Electronics and Computer Technology Center (NECTEC), National Science and Technology Development Agency (NSTDA), Khlong Luang, Pathumthani 12120, Thailand; Department of Chemistry, The University of Western Ontario, 1151 Richmond Street, London, Ontario N6A 3K7, Canada; Department of Applied Mathematics, The University of Western Ontario, 1151 Richmond Street, London, Ontario N6A 5B7, Canada; The Centre for Advanced Materials Research (CAMBR), The University of Western Ontario, 1151 Richmond Street, London, Ontario N6A 5B7, Canada

## Abstract

Carbon nanoparticles (CNPs) are attractive materials for a great number of applications but there are serious concerns regarding their influence on health and environment. Here, our focus is on the behavior of fullerenes in lipid bilayers with varying lipid saturations, chain lengths and fullerene concentrations using coarse-grained molecular dynamics (CG-MD) simulations. Our findings show that the lipid saturation level is a key factor in determining how fullerenes behave and where the fullerenes are located inside a lipid bilayer. In saturated and monounsaturated bilayers fullerenes aggregated and formed clusters with some of them showing icosahedral structures. In polyunsaturated lipid bilayers, no such structures were observed: In polyunsaturated lipid bilayers at high fullerene concentrations, connected percolation-like networks of fullerenes spanning the whole lateral area emerged at the bilayer center. In other systems only separate isolated aggregates were observed. The effects of fullerenes on lipid bilayers depend strongly on fullerene aggregation. When fullerenes aggregate, their interactions with the lipid tails change.

## Introduction

Carbon nanoparticles (CNPs) have been extensively used in various applications such as nanoelectronics, energy, optics cosmetics and nanomedicine [1-4]; particular examples include drug delivery, bioimaging and biosensing [3, 5]. The amount of fullerenes in the environment is also increasing as a result of industrial processes and emission from natural sources, and they are also released as combustion products of fuel engines and heating plants [6, 7]. Thus, the possible health and environmental effects of fullerenes are a growing concern. [8-11]

Previous studies of fullerenes on the effects of fullerenes on biological activities have revealed both positive and negative impacts depending upon many factors. [12-17] The uptake of C60 by human macrophage cells and the resulting aggregation within the cytoplasm, lysosomes and cell nuclei have been visualized using electron microscopy. [18] Additionally, it is remarkable that fullerenes can enter through outer cell membranes, and in some cases, penetrate into the cellular nuclei. [19-21] Although many studies [22, 23] have demonstrated that fullerenes at low concentrations have no significant effects on the physical properties of lipid bilayers, the question whether pristine fullerene physically damages lipid bilayers remains.

A number of studies [22, 24] show that fullerenes neither aggregate inside lipid bilayers nor cause physical damage. On the other hand, at high concentrations several experimental and computational studies have shown that fullerenes aggregate and may be harmful to biological membranes [24-31]. Even the deceivingly simple question of whether fullerenes aggregate inside membranes remains controversial. Considering the fact that the membrane thickness is of the order of a few nanometers and the average distance between neighboring lipid molecules is in the subnanometer length scale, the aggregation mechanisms inside membranes cannot be easily investigated. Despite obvious practical interest, no dedicated studies focusing on the interactions of fullerenes with realistic models of eukaryotic cell membranes have been performed to-date.

To the best of our knowledge, most computational studies of fullerenes in lipid membranes deal with single component lipid bilayers. Our previous computational studies [24] show that fullerene aggregation and its physical mechanisms depend on the degree of lipid unsaturation. Open questions include if the position(s) and the number of double bonds are important in fullerene-lipid interactions. Such knowledge is a necessary pre-requisite for the development of better predictive models for real biological membranes and studies of more realistic multicomponent membranes.

In this study, we performed CG-MD simulations to investigate the physical mechanisms of fullerene aggregation by varying the concentration of fullerenes and the types of lipid bilayers. The results show the dispersion and aggregation of fullerenes depends both on degree and position(s) of lipid unsaturation(s). Aggregation were analyzed to explain the potentially harmful effects on biological membranes. This provides knowledge of which features of fullerene aggregation inside membranes determine toxic responses, as well as help with the identification of the corresponding toxicity.

## Methodology

### MD simulation

We performed CGMD simulations to study fullerene behavior inside lipid bilayers and the effect of fullerenes on lipid membranes. Seven types of phospholipids displaying different number double bonds and tail lengths were used, Figure 1A and S1: 1,2-dilauroyl-sn-glycero-3-phosphocholine (DLPC), 1,2-dipalmitoyl-sn-glycero-3-phosphocholine (DPPC), 1,2-distearoyl-sn-glycero-3-phosphocholine (DSPC), 1,2-dioleoyl-sn-glycero-3-phosphocholine (DOPC), 1,2-divaleryl-sn-glysero-3-phosphatidylcholine (DVPC), 1,2-diundecanoyl-sn-glycero-3-phosphocholine (DUPC) and 1,2-dioctadecatrienoyl-sn-glycero-3-phosphocholine (DFPC). Of the above, DLPC (12-14:0), DPPC (16-18:0) and DSPC (20-22:0) are saturated with different chain lengths, whereas DOPC (16-18:1), DVPC (16-18:1), DUPC (16-18:2) and DFPC (16-18:3) are unsaturated with different number of double bonds. Although DOPC and DVPC have the same chain lengths and number of double bonds, the locations of double bond in DOPC and DVPC are different. The MARTINI force field version 2.1[32, 33] was applied for lipids and water. The coarse-grained fullerene model (F16) which represents the C60 molecule was taken from the latest version [34]. The fullerene model consists of 16 beads and reproduces the atomistic potential of mean force (PMF) profile of transferring a fullerene across lipid membrane. [34] The systems contained 512 lipid molecules and 16,000 CG water beads. Fullerenes were randomly placed above the lipid bilayers with fullerene/lipid ratios of 0%, 5%, 10%, 20%, 30% and 40%. The details of all simulations are shown in Table S1. Simulations were performed under the NPT ensemble using the GROMACS 4.5.5 package [35]. The temperature was kept at 298 K using the Parrinello–Donadio–Bussi velocity rescale algorithm [36, 37] with a time constant of 1.0 ps. The Parrinello–Rahman algorithm [38] with semi-isotropic coupling was used for keeping pressure constant at 1 bar with a time constant 5.0 ps and compressibility of 4.5 × 10^−5^ bar^−1^. A cut-off distance of 1.2 nm was employed for non-bonded interactions. The Lennard-Jones interactions were shifted to zero between 0.9 and 1.2 nm and the Coulomb potential was shifted to zero between 0 and 1.2 nm. To remove possible molecular overlaps that may have occurred during the setup, the systems were first subjected to energy minimization using the steepest descent method. The simulations were run for 20-40 *μ*s with a 20 fs time step. The last 10 *μ*s were considered for analysis. Periodic boundary conditions were applied in all three dimensions. The Visual Molecular Dynamics (VMD) software was used for rendering molecules [39]. The MDAnalysis [40, 41] and NetworkX [42] python libraries were used for analysis.

**Figure 1.**
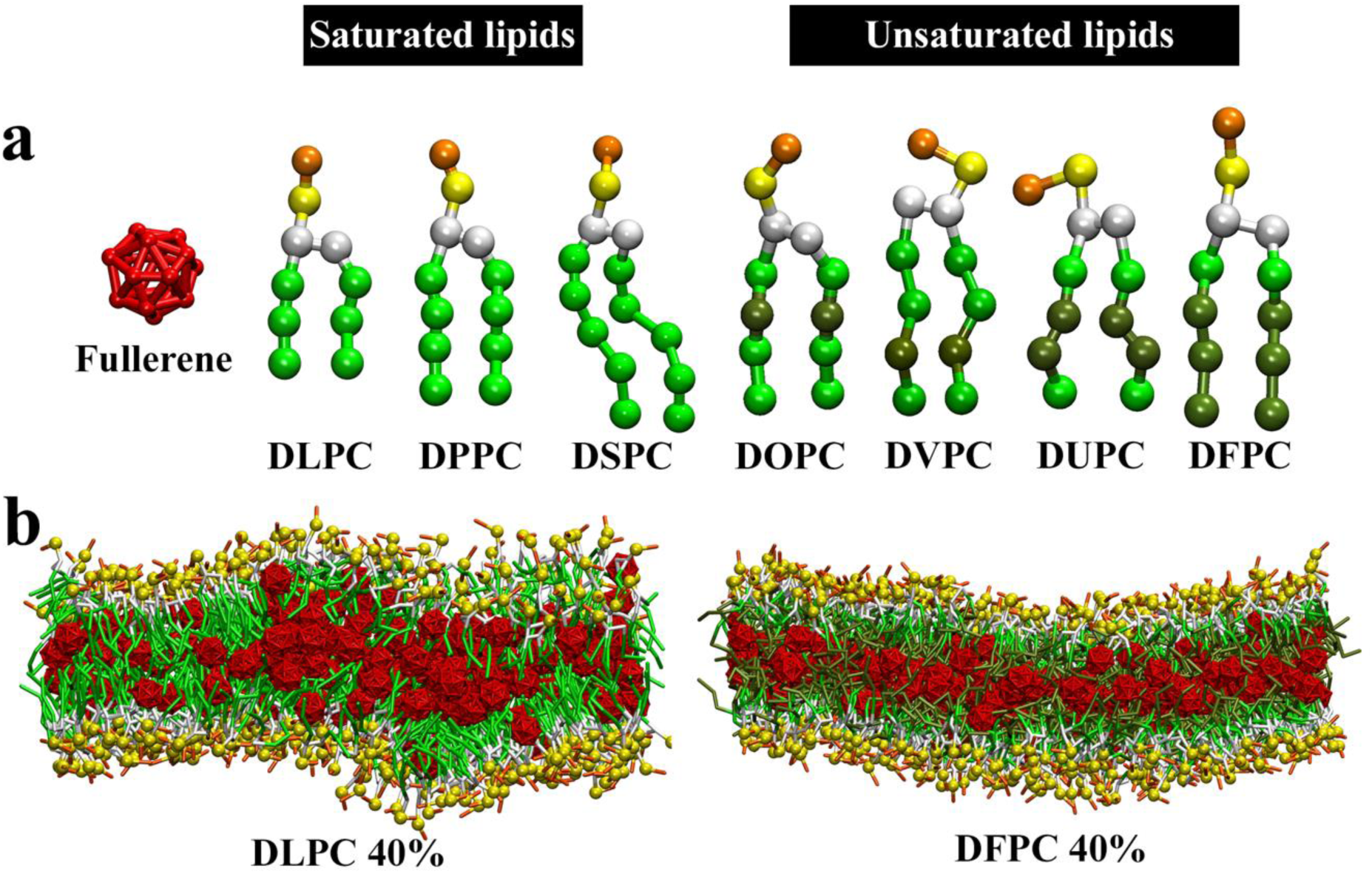
(a) The MARTINI coarse-grained (CG) models of fullerene and various lipid types. Green corresponds to apolar beads in the lipid tails (normal: saturated carbons, dark: double bonds). Red, orange, yellow and white beads represent fullerene, choline, phosphate and glycerol groups, respectively. Saturated lipids are DLPC (12-14:0), DPPC (16-18:0), and DSPC (20-22:0). Unsaturated lipids are DOPC (16-18:1), DVPC (16-18:1), DUPC (16-18:2) and DFPC (16-18:3). (b) DLPC and DFPC bilayers with fullerene concentration of 40% after 20 microseconds. Water molecules are not shown for clarity.

### Potential of mean force (PMF)

To calculate the PMF for a single fullerene transferring through a lipid bilayer, the umbrella sampling technique with the Weighted Histogram Analysis Method (WHAM) [43, 44] was applied. DLPC, DSPC and DFPC bilayers at two fullerene concentrations (0% and 40%) were studied. The initial structures were taken from the last frames of the unbiased simulations. The distance between the center of mass of the fullerene and the bilayer was restrained in the z-direction with a harmonic potential force constant 1000 kJ mol^−1^. A single fullerene was initially placed in the water phase (at z=4.5 nm) and pulled to the center of the bilayer (z=0 nm) with 0.1 nm increment. All simulations were performed under the NVT ensemble at 298 K for the total time of 46 *μ*s (1 *μ*s per each window). Error analysis was done with the bootstrapping analysis method [44] for complete histograms.

## Results

### Fullerenes’ locations inside the bilayers

Previous studies of fullerene permeation mechanisms into bilayers have established that the energy barrier for a fullerene entering into a lipid bilayer is negligible (i.e. less than *k*_*B*_T). [22, 45] Our simulations show that once fullerenes enter a bilayer, they stay inside it as seen in Figures 1, S2 and S3. Interestingly, our simulations also show that the final location of fullerenes inside a bilayer depends strongly on fullerene concentration and lipid type, Figure 2. In saturated bilayers (DLPC, DPPC, and DSPC) and at low fullerene concentrations (i.e. less than 20%), fullerenes prefer to be located in the hydrophobic acyl chain region but away from the bilayer center. The potential of mean force (PMF) calculations in Figure 3 shows that the free energy minima from the bilayer center were ∼0.7, ∼1.1, and ∼1.4 nm for the DLPC, DPPC and DSPC bilayers, respectively. The location of the free energy minimum in the DLPC bilayer agrees with the all-atom PMF study of C60 in DMPC bilayers; [46] note that CG-DLPC (12-14:0) is equivalent to all-atom-DMPC (14:0). When increasing fullerene concentration to 30% and 40%, the fullerene density separates into 3 regions including the bilayer center, Figure 2. Unlike for other saturated lipids, for the shortest of them (DLPC) fullerene density at the bilayer center reached saturation at about 557 kg m^−3^ at 40% fullerene concentration. Note that saturation density of about 547-557 kg m^−3^ was observed in unsaturated DVPC and DUPC lipid bilayers at 30% and 40% fullerene concentrations.

**Figure 2.**
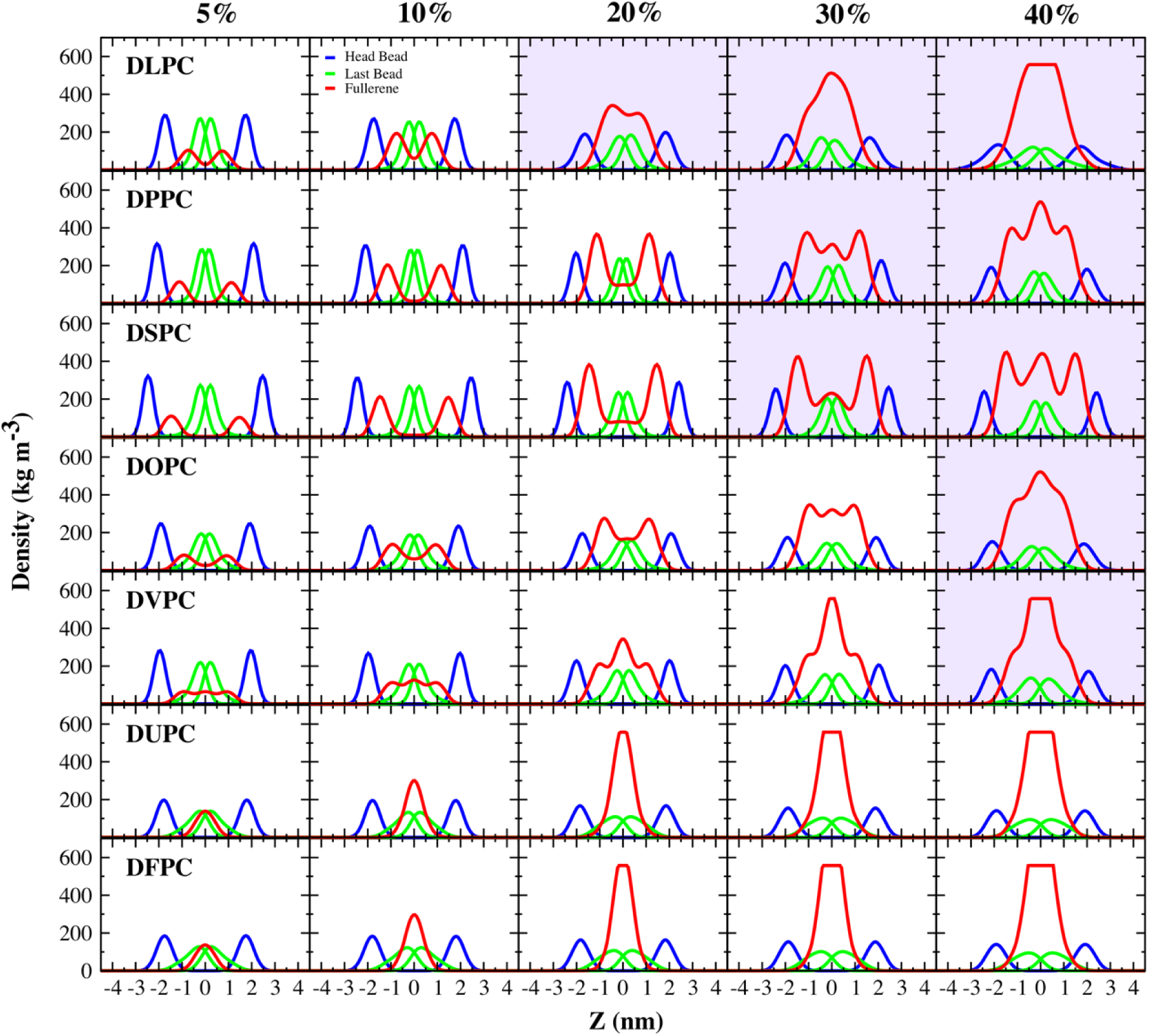
Bilayer density profiles at different fullerene concentrations along the bilayer normal (z-axis). All profiles shown are symmetrized over the two leaflets of the bilayer. Note that the lipid head bead is PO4 and the last beads are C3A (DLPC), C4A (DPPC, DOPC,DVPC,DUPC), C5A (DSPC) and D4A (DFPC). The structures and lipid indices are provided in Figure S1. Purple background color represents fullerene aggregation.

**Figure 3.**
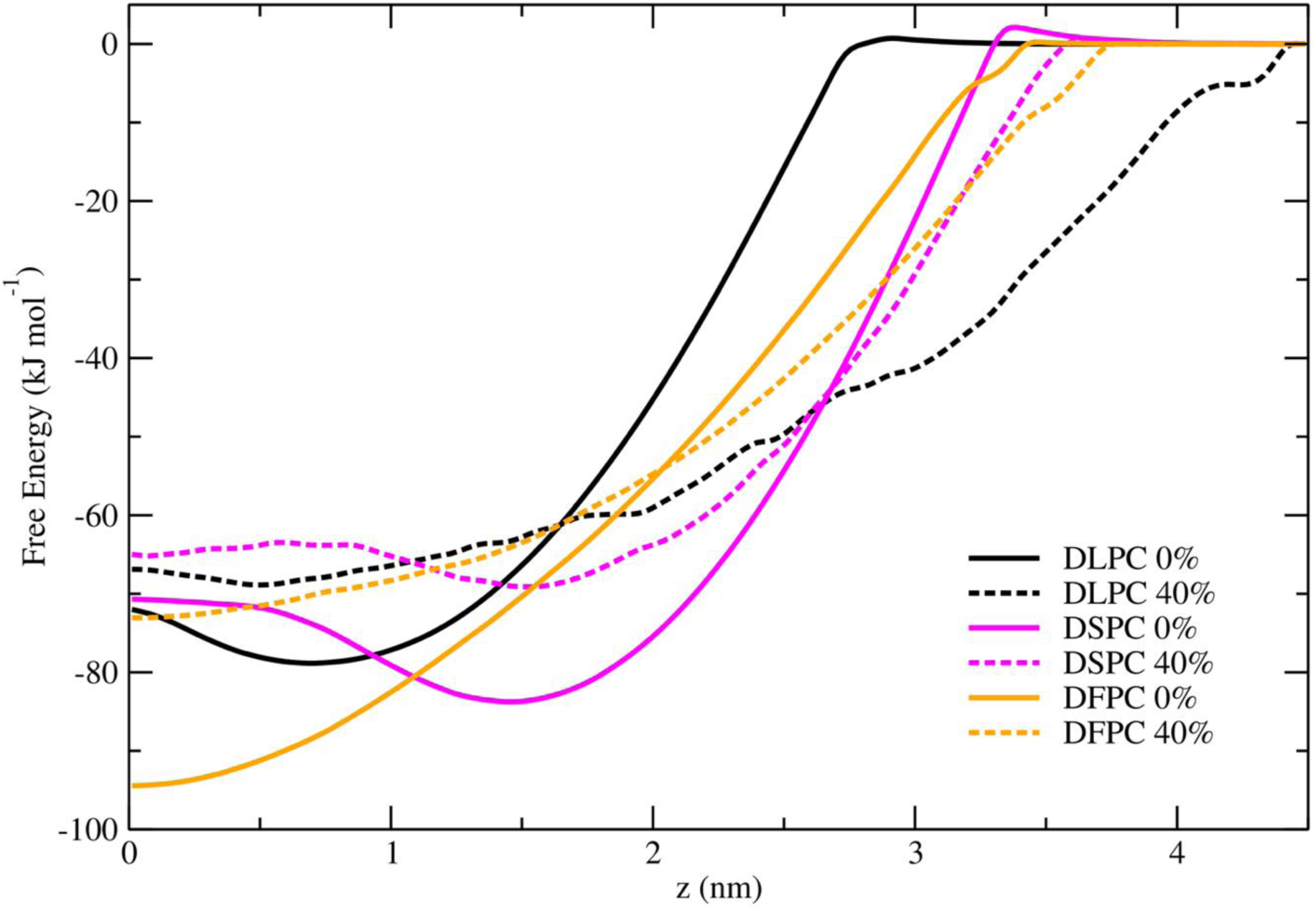
Potential of mean force for fullerene translocation across the bilayer from the water phase into the center of the lipid bilayer (z=0 nm) as a function of distance in the z-direction. The solid and dashed lines correspond to fullerene concentrations of 0% and 40%, respectively. The estimated errors are less than 0.887, 1.902, 0.430, 1.217, 1.651, and 0.930 kJ mol^−1^ for DLPC 0%, DLPC 40%, DSPC 0%, DSPC 40%, DFPC 0%, and DFPC 40%.

Different fullerene behavior was observed in unsaturated lipid bilayers. Independent of concentration, fullerenes in unsaturated bilayer preferred to be located at center of the bilayer to avoid contacts with the double bond regions. These observations are corroborated by our PMF calculations, Figure 3.

In both saturated and unsaturated bilayers, fullerenes became less stable inside the bilayer as the concentration was increased. One possible explanation of this behavior is the decrease in free volume upon increasing fullerene density. Interestingly, the penetration barrier free energies for fullerenes into the DFPC bilayer were extremely low (i.e. close to zero). When comparing the free energy minimum for fullerene penetration in the cases of pure DFPC lipid bilayer and 40% of fullerene, the former one is the most stable case with the minimum free energy of −94.4 ± 1.6 kJ mol^−1^ (Figure 3). In addition, the wrinkly shape of the free energy profile for the DLPC bilayer at 40% fullerene concentration is a result of perturbation by fullerenes (Figure 3).

### Fullerene aggregation inside lipid bilayers

The contradiction whether fullerenes form cluster(s) inside lipid bilayer has been discussed for a decade. In 2007, Porter et al. demonstrated that fullerenes (C60) aggregate in plasma membranes by using electron microscopy techniques.[18] However, the recent study of Zhou et al. [47] used atomic force microscope (AFM), laser light scattering (LLS) and Fourier-transform infrared spectroscopy (FTIR), and found that fullerenes dispersed in saturated bilayer. The explanation behind these observations is unclear as used different techniques and bilayer types were used. The situation is even more complicated when looking at computational studies. Current studies support both aggregation and dispersion of fullerenes in bilayer depending on fullerene concentration, fullerene cluster size in aqueous solution and lipid bilayer composition.[22-24, 28, 31, 46, 48-55] To-date, there was no systematic study investigating this mysterious.

To improve our understanding of fullerene aggregation and dispersion, we performed a series of systematic analyses using seven different bilayers and five fullerene concentrations. We first calculated the C60-C60 center of mass radial distribution function (RDF), Figure 4. At 10% fullerene concentration, the RDFs are similar in all systems and in agreement with previous studies.[24, 56] There are two peaks at about 1 nm and 1.5 nm. When increasing fullerene concentration, the difference in the RDFs of fullerenes in saturated and polyunsaturated bilayer becomes observable. In saturated and monounsaturated bilayers, an additional peak occurring about 1.65 nm was observed at the fullerene concentrations higher than 10%. Furthermore, the peaks at about 2 nm became more pronounced. These peaks represent the optimized distances as demonstrated in Figure 4. The first peak (at ∼1 nm) is the distance between the centers of masses of two fullerenes forming a dimer. It also represents the distance for the nearest neighbors, or first shell, when fullerenes form a cluster. The second peak (at ∼1.5 nm) is the configuration when a lipid tail has become inserted between two fullerenes. These two characteristics have been already shown and verified in studies of C60 in ethanol solution [57] and in membranes.[24] The third peak (at ∼1.65 nm), according to Ding et al. [58], is the splitting peak of the second peak that implies cluster connections (i.e. second shell). The fourth peak (at ∼2nm) represents the third shell.

**Figure 4.**
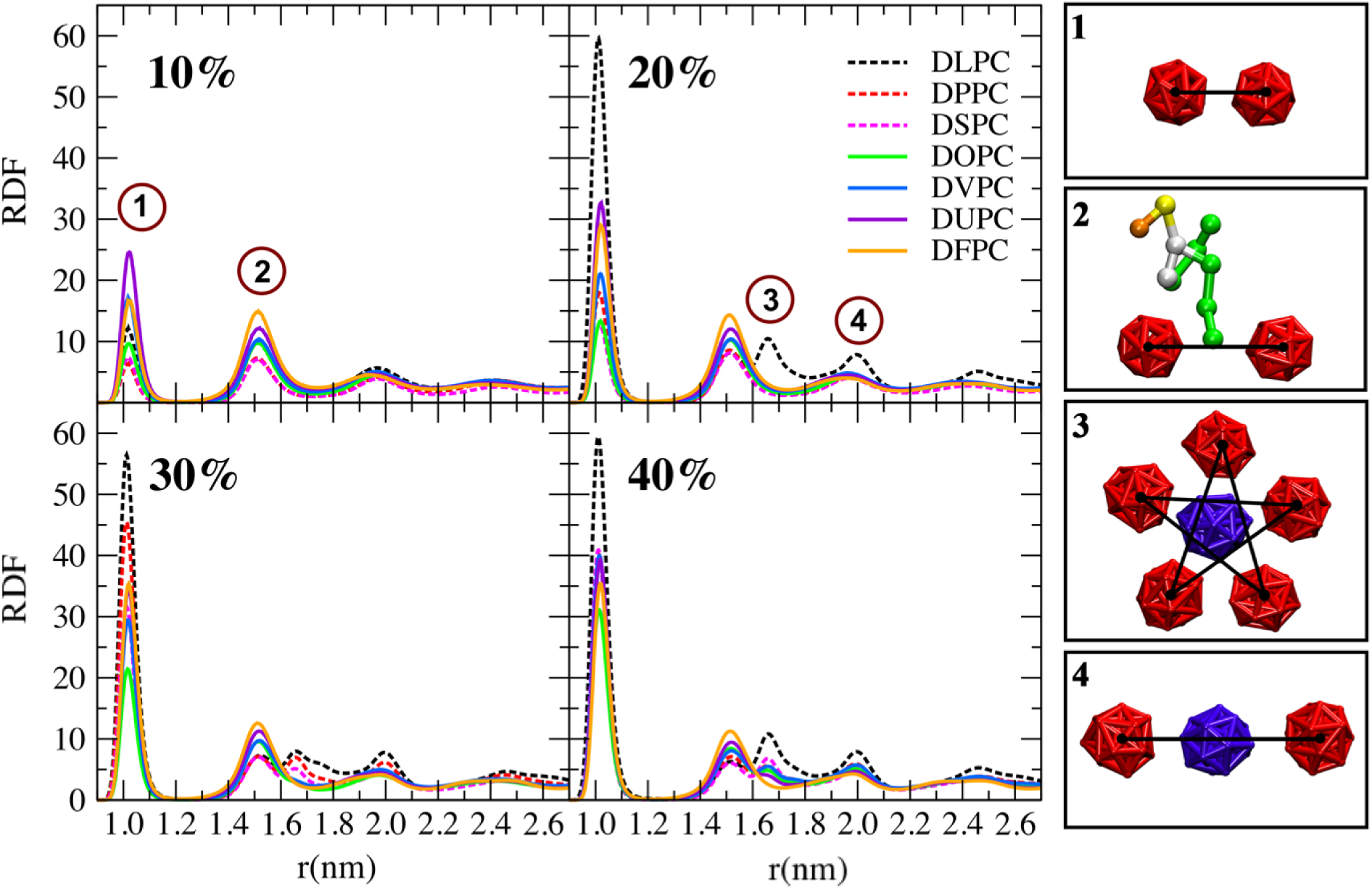
Fullerene-fullerene radial distribution functions (RDFs) inside the lipid bilayers at different fullerene concentrations (left). Snapshots of representative configurations corresponding to the numbers (right). An additional peak of 1.65 nm was observed at fullerene concentrations higher than 10%.

It was observed that the C60 molecules formed the so-called Mackay’s icosahedral structure [59] as illustrated in Figure 5. It is a stack of AB layers where layer A has five-fold symmetry and is formed by five C60 molecules in the same plane and layer B is a single C60 molecule. Because of lipid tails, some of the clusters observed in the simulations were not perfect. Instead of 12 neighbor molecules (for (C60)_13_ clusters), there were atoms missing in some positions. Icosahedral structures of fullerenes were first found using photoionization time-of-flight mass spectrometry technique.[60] In addition, they have also been predicted by theoretical approaches.[61-65] In a recent CG-MD simulation study of Xie et al., [48] icosahedral fullerene structure was observed in water phase and the five-fold symmetry plane orientations were random as can be expected by the rotational and translational symmetries of the water phase. In contrast, for fullerenes to be able to cluster inside a bilayer, the five-fold symmetry planes must be aligned with bilayer plane. To ensure that our observations of Mackay icosahedra were not caused by the semi-isotropic pressure coupling that is standard in bilayer simulations, we performed simulations of C60 in water using two different pressure couplings (semi-isotropic and isotropic), the details are provided in SI. In both cases, we found that the C60 molecules formed a large cluster with an icosahedral structure. Importantly, the five-fold symmetry planes were not aligned with the z-axis confirming their independence from the pressure coupling method, see Figure S4.

**Figure 5.**
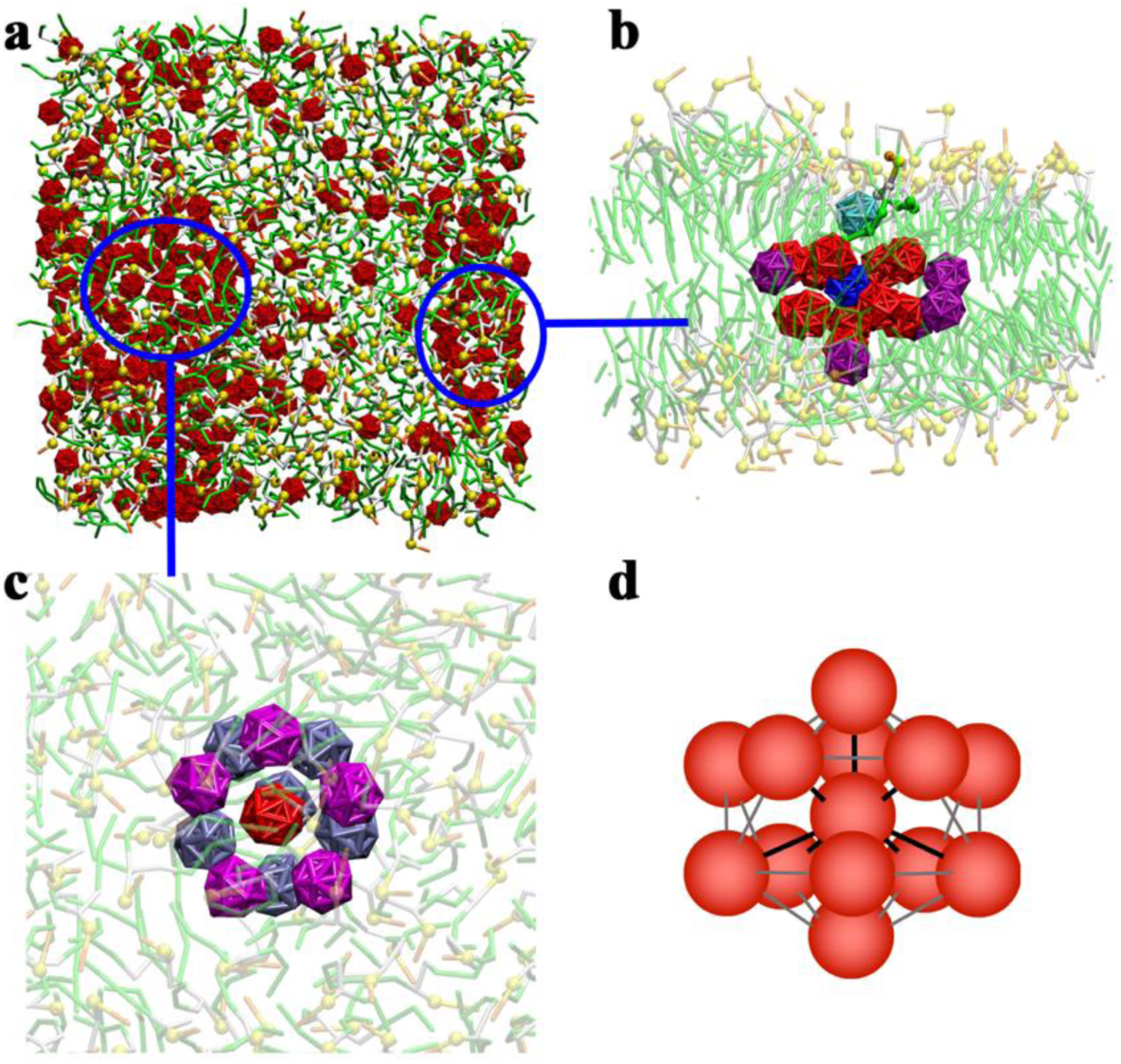
(a) Top view of a DLPC lipid bilayer at 40% fullerene concentration. (b) Side view of the indicated area showing an approximate icosahedral structure caused by lipid tail insertion. (c) Top view of the indicated area showing C60s form an icosahedral cluster and (d) a schematic diagram of an icosahedral cluster. The Mackay icosahedron is a stack of AB layers where the layer A has five-fold symmetry and is formed by five C60 molecules in the same plane and layer B is a single C60 molecule.

### Fullerene clustering in lipid bilayers

The coordination numbers of fullerenes in lipid bilayers were determined by the summation of fullerene molecules within the first shell (a cut-off of 1.3 nm). The results are shown in Table S2. The largest coordination number is at 40% fullerene concentration in the DLPC bilayer. Figure 6 shows the distributions of coordination numbers at 40% concentration in the DLPC and DFPC bilayers. Figure 6a shows that the probability remained close to a constant up the coordination number of 11. On the other hand, in the case of DFPC (Figure 6b) the behavior is qualitatively different with a peak at lower numbers of two and three. Geometrically, the coordination number of two corresponds to (linear) pairs and three to planar trigonal structures. The visualizations in Figures 6e, 6f and Figure S2 also show that the fullerenes are laterally spread in the mid-plane at the DFPC bilayer center. Unlike fullerenes in water, the fullerenes in this plane do not minimize their surface because they are located in a hydrophobic environment. This behavior was also found in other polyunsaturated bilayer systems (Figure S2). Furthermore, as Figure 7a shows, in DLPC the fullerenes form a localized aggregate whereas in DFPC (Figure 7b) the behavior is qualitatively different and a percolation-type cluster emerges.

**Figure 6.**
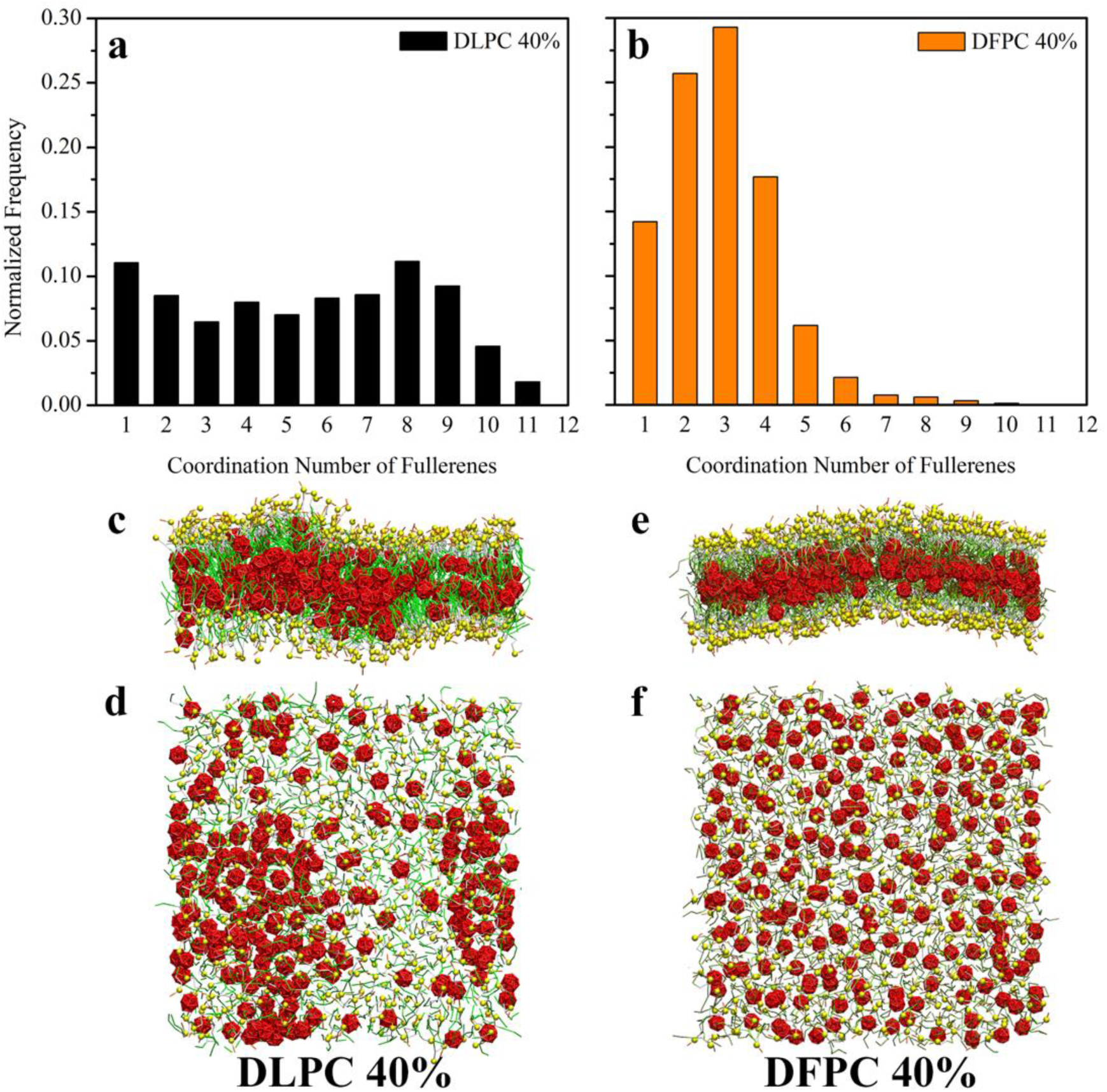
The distribution of fullerene coordination numbers within the cut-off of 1.3 nm in (a) DLPC and (b) DFPC bilayers at 40% fullerene concentration. Side and top views of the DLPC (c-d) and DFPC (e-f) bilayers at 40% fullerene concentration.

**Figure 7.**
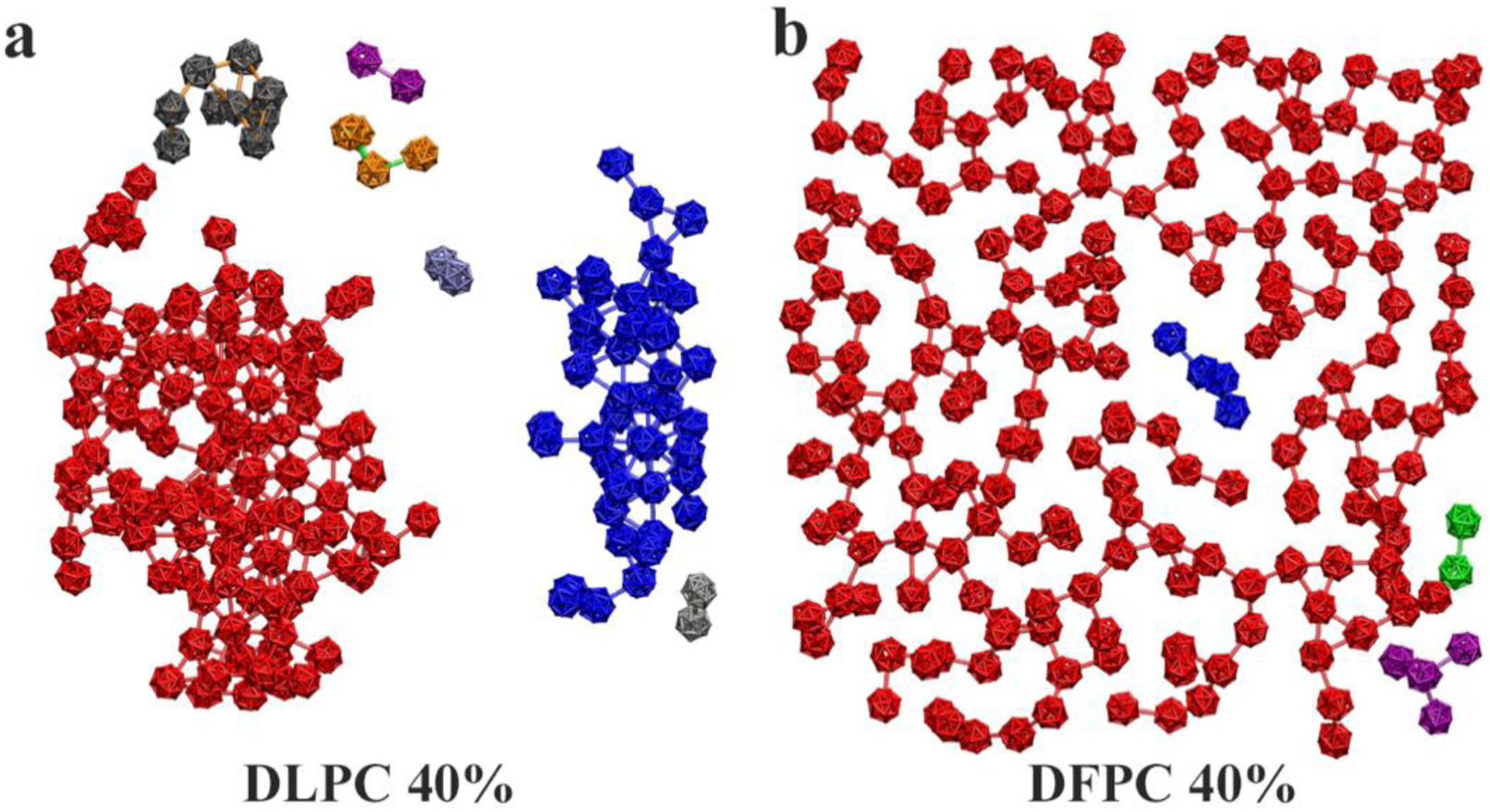
Connected networks of fullerenes within 1.3 nm cut-off (connecting lines). The different clusters are shown using different colors. In DLPC, aggregation to densely packed clusters occurs whereas in DFPC, a qualitatively different a percolation-like network emerges.

### Fullerene diffusion

We calculated diffusion coefficients using mean square displacements and Einstein’s relation. In agreement with other studies [24, 26, 31], fullerenes moved slower at high concentration, see Figure 8. Interestingly, we observed clear differences between saturated and highly unsaturated lipid bilayers: fullerenes in saturated lipid bilayer move slower than in unsaturated ones. A possible explanation is the different locations of the fullerenes in these bilayers. In saturated bilayer, fullerenes located in the acyl chain region and therefore their movement is restricted. On the other hand, in unsaturated bilayers, fullerenes are located in the middle of bilayer which has more free space. This difference was very small at high concentration because the motion of fullerenes becomes limited due to crowding. Lateral fullerene diffusion in DVPC, DOPC and DLPC bilayers is quite similar because the differences in their structures are minor. This explains why in our previous work [24] did not observe differences between DPPC, POPC, and DOPC bilayers.

**Figure 8.**
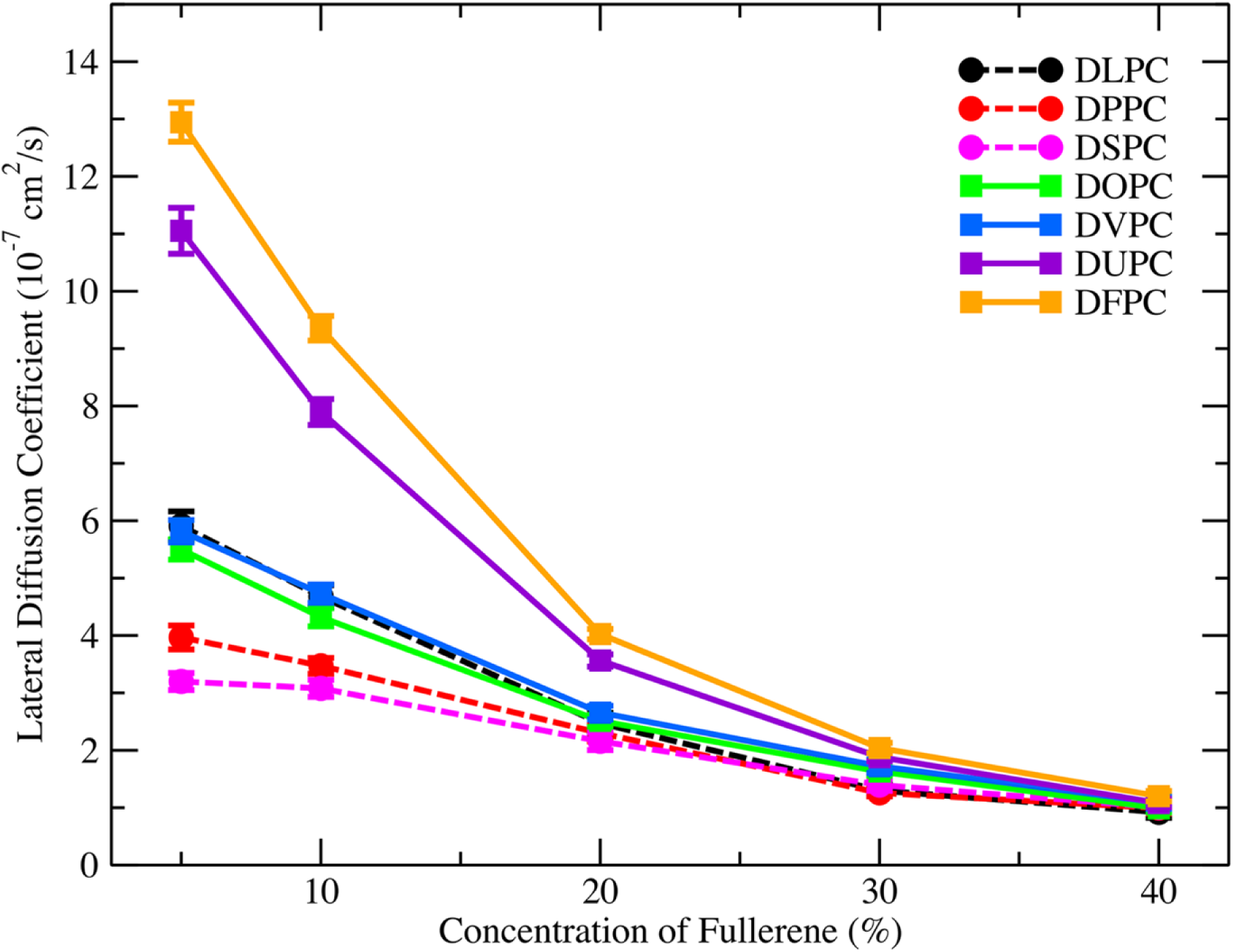
Fullerene diffusion coefficients as a function of fullerene concentration

### Effect of fullerenes on bilayers

To study the physical and mechanical properties of bilayer in the presences of fullerenes, we computed the area per lipid, bilayer thickness, and volume per lipid of the bilayer as seen in Table S3 and Figure S5. In the absence of fullerenes or at low fullerene concentration, our results are in agreement with previous studies [22-24, 28, 31, 48, 49], as show in Figure 9 and S5. Differences emerged when fullerene concentration was increased. In general, area per lipid, bilayer thickness, and volume per lipid increase with increasing fullerene concentration. However, the thickness of saturated bilayers decreases at low fullerene concentrations and then increases at high fullerene concentration. These changes were also observed in our previous study [24]. A possible explanation is that individual fullerenes and fullerene clusters behave differently in these situations since they started to aggregate at high concentration in saturated bilayer: (10% for DLPC, 20% for DPPC, 20% for DSPC).

**Figure 9.**
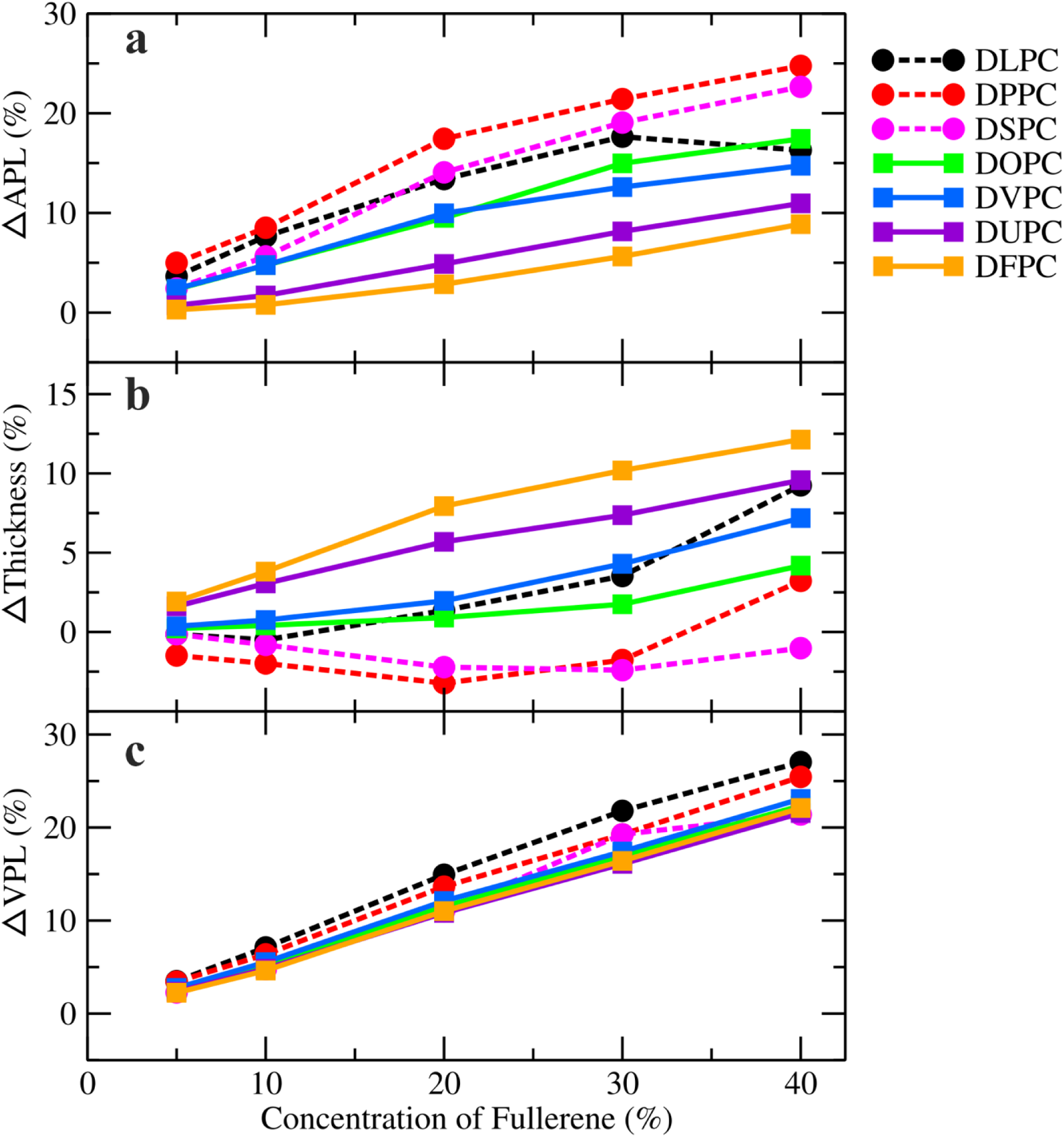
(a) Area per lipid, (b) bilayer thickness and (c) volume per lipid as a function of fullerene concentration in the different lipid bilayers. The dashed and solid lines correspond to saturated and unsaturated lipid bilayers, respectively. Error bars are smaller than the size of the symbols.

To understand how fullerene aggregation changes bilayer properties, we analyzed local properties. First, we calculated the lipid tilt angles, that is the angle between the z-axis and the vector connecting the PO4 and the different tail beads as shown Figure 10. The results show that tail orientations of DPPC and DFPC lipids are distinct. In case of DPPC at low concentrations (≤20%), fullerenes increase the tilt angle of C1, but angles of C2, C3, and C4 remain constant or decrease slightly (all defined using PO4). At high concentrations (>20%) also the C2, C3 and C4 angles increase with increasing fullerene concentration.

**Figure 10.**
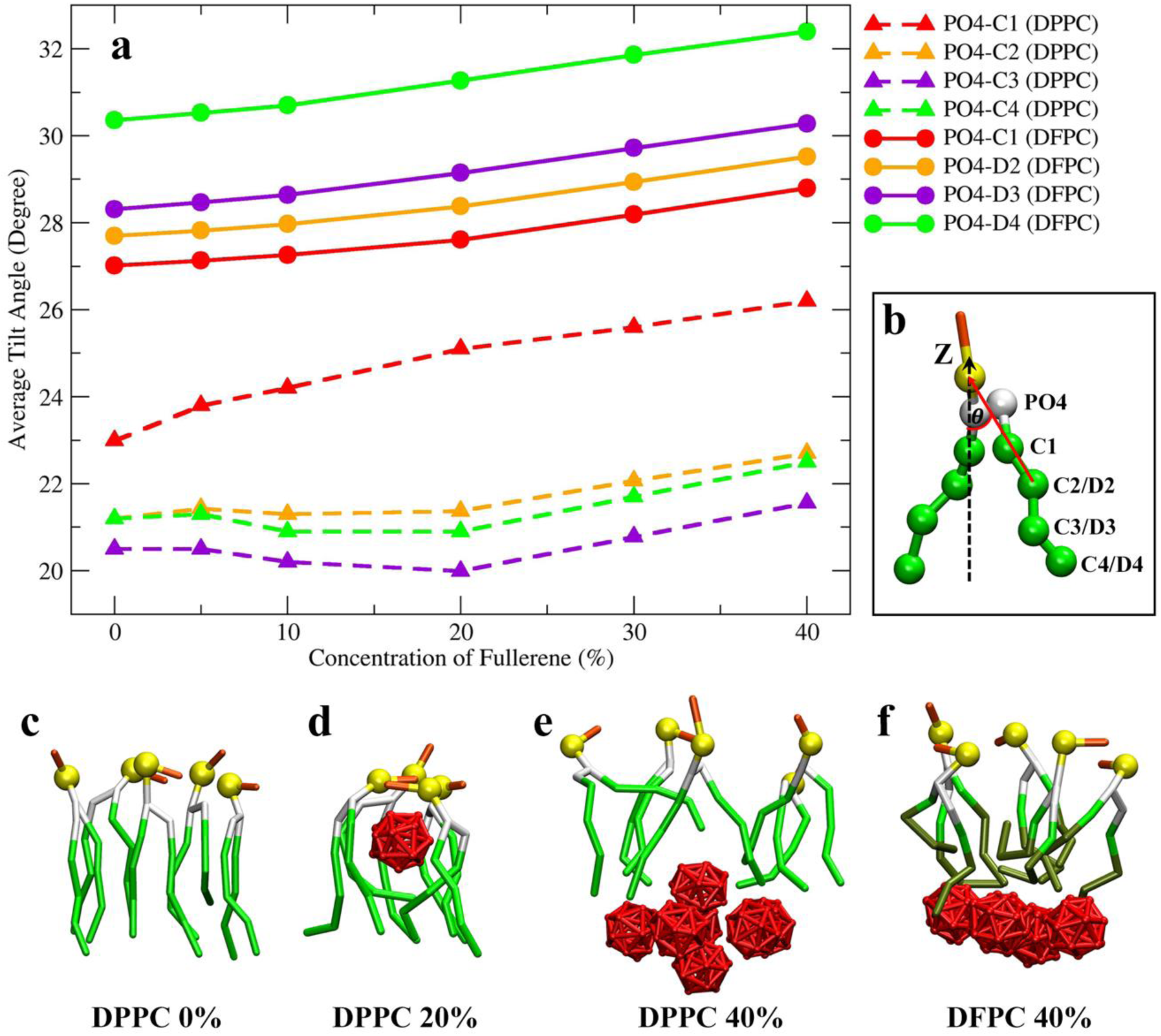
**a)** The average tilt angles of each lipid tails in the DPPC and DFPC bilayer at different fullerene concentrations computed as shown in b). Dash lines: DPPC bilayer. Solid lines: DFPC bilayer. c-e) Snapshots of DPPC lipids at fullerene concentration of 0%, 20%, 40%, respectively and f) snapshot of DFPC lipids at fullerene concentration of 40%.

In case of DFPC, the tilt angles increase with increasing fullerene concentration (Figure 9). These changes explain how fullerene concentration is related to bilayer thickness. In saturated bilayer, fullerenes are located in the acyl chain region and cause a larger tilt therefore inducing a decrease in bilayer thickness. At high concentrations, fullerene aggregation changes lipid tail orientation and increases bilayer thickness because the size of aggregated fullerene is larger than the excess free volume. In DFPC, fullerenes locate in the middle of bilayer resulting in stretching of the lipid tails and creation of free space for fullerenes.

In addition, we analyzed the local 2D density and thickness. Figures 11, S6 and S7 demonstrate that local increases in bilayer thickness are directly correlated with the locations of fullerene clusters; the area per lipid increases monotonously with fullerene concentration. We would like to point out that the extensive drop in DLPC area per lipid at 40% is caused by bilayer bending. These results reveal that fullerenes’ effects depend on many factors: saturation level, double bond positions and acyl chain length. Importantly, the differences between saturated and unsaturated are clearly observable. This emphasizes the significance of lipid saturation level since fullerenes behave very differently in saturated bilayer and unsaturated bilayers.

**Figure 11.**
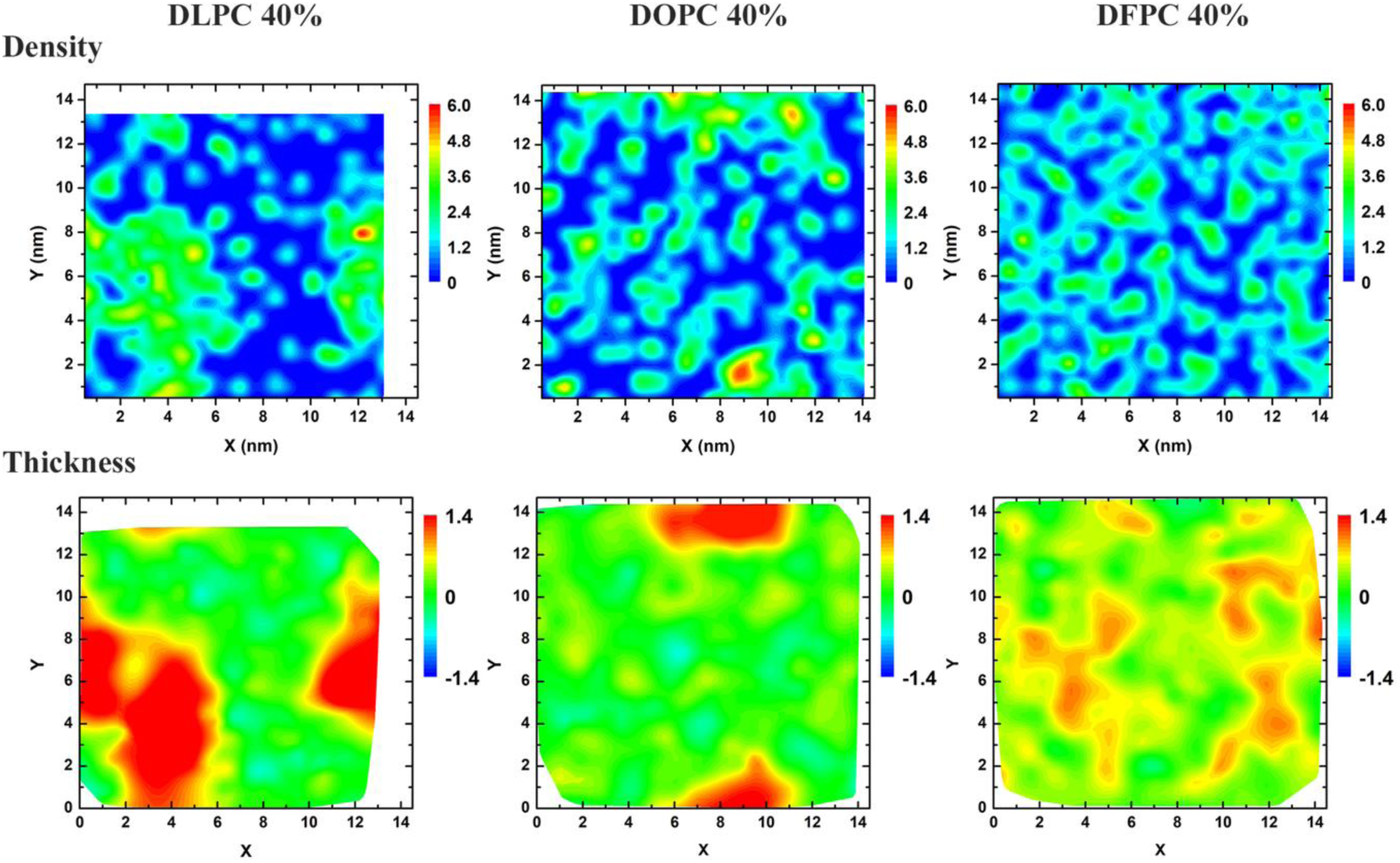
2D fullerene density map and bilayer thickness at 40% fullerene concentration in DLPC, DOPC and DFPC lipid bilayers. The increase in bilayer thickness is correlated with the locations of the fullerene clusters.

## Discussion

Our systematic study reveals that fullerenes behavior in lipid bilayers depends strongly on i) fullerene concentration, ii) the saturation level in terms of both positions and number of double bonds in the lipid tails and, iii) the acyl chain length. First, the fullerene concentration must be high for aggregation to occur. Second, the level of hydrocarbon chain saturation plays an important role in fullerene clustering. When increasing fullerene concentration, the differences in fullerenes’ behaviors in these bilayers becomes more pronounced. Some of the mechanisms behind these observation were already explained in our previous study [24] which discovered that the balance between entropy and enthalpy is controlled by fullerene aggregation. In saturated bilayers, the entropic component decreases faster than the enthalpic component increases, whereas, in unsaturated bilayer, the entropic component dominates the free energy. Third, acyl chain length also plays role in fullerene aggregation. It determines the amount of fullerene absorbed by bilayer before cluster formation or reaching (fullerene) saturation level. For saturated (DLPC, DPPC, DSPC) and unsaturated bilayers (DOPC, DVPC), the fullerene saturation level depends directly on acyl chain length as the fullerenes are located in the acyl chain region. For polyunsaturated bilayers (DUPC, DFPC), the longer lipid tails also mean that the lipids have higher adjustability for larger free space in bilayer center. Based on our DOPC and DVPC simulations, we found that role of position of double bond is minor.

In terms of toxicity as determined by membrane damage, we did not observe physical membrane damage or rupture during any of the simulations. However, fullerene aggregation strongly disturbs lipid bilayers resulting in bending as shown in some of the snapshots at high concentrations in Figure S2. In addition, the biological consequences of localized aggregation (Figure 7a) and the formation of percolation clusters (Figure 7b) are currently not known. Moreover, there are experimental [47] and computational [26] studies that have demonstrated that fullerenes could significantly modify bilayers’ mechanical properties. In addition, an increase in cluster size in aqueous solution may induce membrane rupture as it may exceed bilayer thickness [66]. Furthermore, at a certain cluster size, lipid bilayer loses its ability to dissolve fullerene [48]. However, Kyzyma et al. [67] did viability tests using living cell cultures and found that the fullerene aggregate size in water appears not to be a key factor for toxicity. Along with this, the experimental study of Fang et al. [68] has also shown that bacteria can adjust themselves as they respond to C60s by altering their membrane lipid compositions. Recent studies also demonstrate interactions between fullerenes and proteins. [69-72] Considering all these factors, determining fullerenes’ toxicity or lack of it on living cells remains a very complex and challenging.

## Acknowledgement

This work was financially supported by Kasetsart University Research and Development Institute (KURDI) and Faculty of Science at Kasetsart University (JW, NN). The Thailand Science Research and Innovation (TSRI), National Research Council of Thailand (NRCT) and Kasetsart University through the research scholarship (Grant No. RSA6180021, PHD/0204/2559, and PHD/0134/2561) to support JW is acknowledged. MK would like to thank the Natural Sciences and Engineering Research Council of Canada (NSERC) and the Canada Research Chairs Program. Computing facilities were provided by SHARCNET (www.sharcnet.ca), Compute Canada (www.computecanada.ca) and the Department of Physics, Faculty of Science, Kasetsart University.

## Supporting Information

### C60s in water simulation

Two simulations were performed using the GROMACS 5.1.1. simulation software [1] After energy minimization, the simulations were performed in the NPT ensemble under two different pressure couplings (semi-isotropic, isotropic). The Parrinello–Rahman algorithm [2] was used to maintain constant pressure at 1 bar with a time constant 12.0 ps and compressibility of 4.5 × 10^−4^ bar^−1^. Temperature was kept constant with the Parrinello–Donadio–Bussi velocity rescale algorithm [3, 4] at 298 K and time constant of τ = 1.0 ps. A cutoff radius of 1.1 nm was applied for the real space part of the electrostatic interactions and Lennard-Jones interactions. Each of the systems contained 256 fullerene molecules and 20,000 CG water beads. The simulations were run for 5 µs with a time step of 20 fs.

**Table S1.** Simulation details

| Fullerene<br>concentration | Molecules |  |  | Simulation time (μs) |  |  |  |  |  |  |
| --- | --- | --- | --- | --- | --- | --- | --- | --- | --- | --- |
|  | Fullerene | Lipid | Water | Saturated lipid bilayer |  |  | Unsaturated lipid bilayer |  |  |  |
|  |  |  |  | DLPC | DPPC | DSPC | DOPC | DVPC | DUPC | DFPC |
| No fullerene | 0 | 512 | 16000 | 20 | 20 | 20 | 20 | 20 | 20 | 20 |
| 5% | 24 | 512 | 16000 | 20 | 20 | 20 | 20 | 20 | 20 | 20 |
| 10% | 48 | 512 | 16000 | 20 | 20 | 20 | 20 | 20 | 20 | 20 |
| 20% | 104 | 512 | 16000 | 30 | 20 | 30 | 20 | 20 | 20 | 20 |
| 30% | 152 | 512 | 16000 | 20 | 30 | 30 | 20 | 30 | 20 | 20 |
| 40% | 204 | 512 | 16000 | 20 | 20 | 40 | 20 | 30 | 20 | 20 |

**Figure S1.**
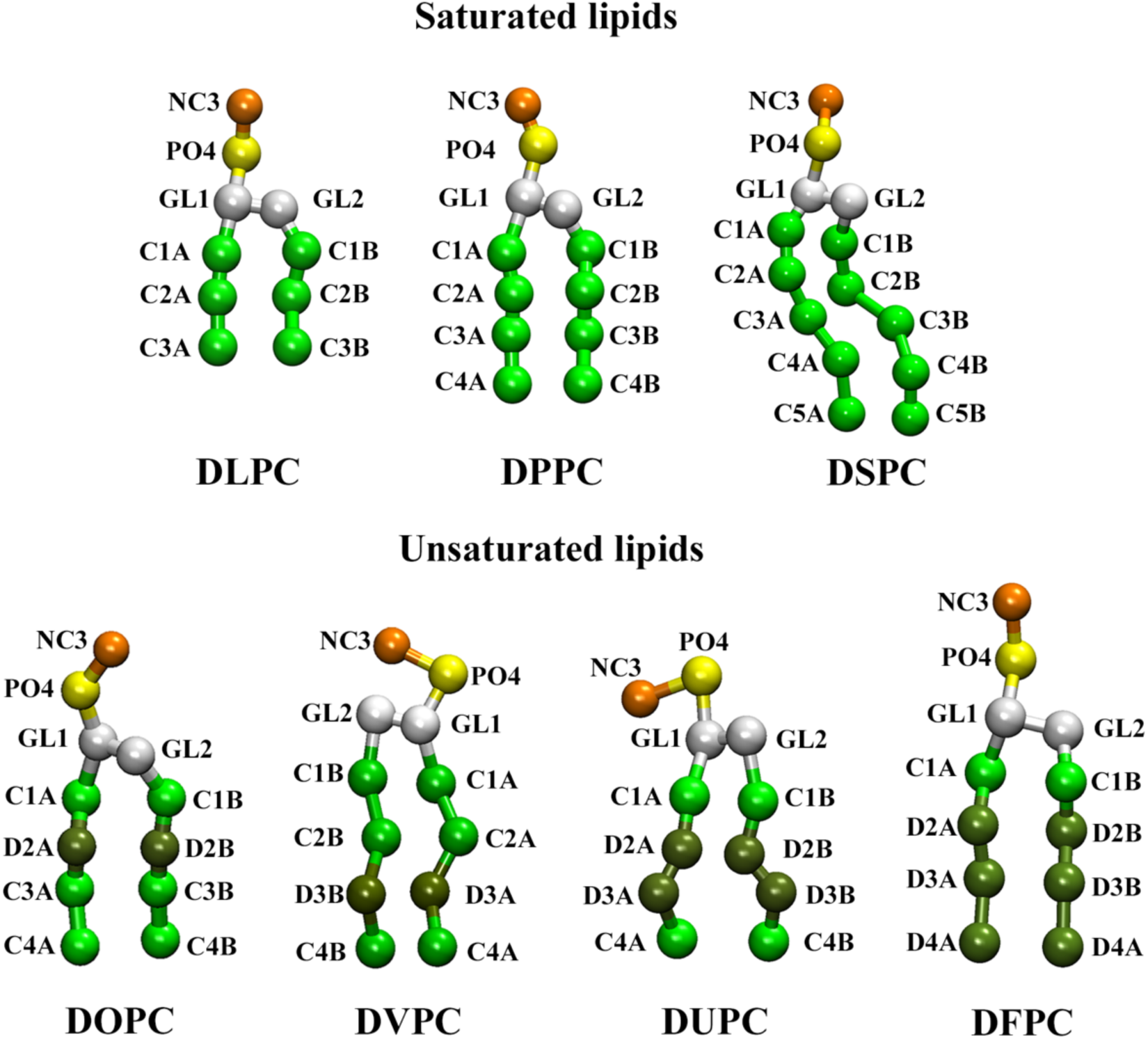
The MARTINI coarse-grained (CG) models of fullerene and various lipid types. Green corresponds to apolar beads in the lipid tails (normal: saturated carbons, dark: double bonds). Red, orange, yellow and white beads represent fullerene, choline, phosphate and glycerol groups, respectively. Saturated lipids are DLPC (12-14:0), DPPC (16-18:0), and DSPC (20-22:0). Unsaturated lipids are DOPC (16-18:1), DVPC (16-18:1), DUPC (16-18:2) and DFPC (16-18:3).

**Figure S2.**
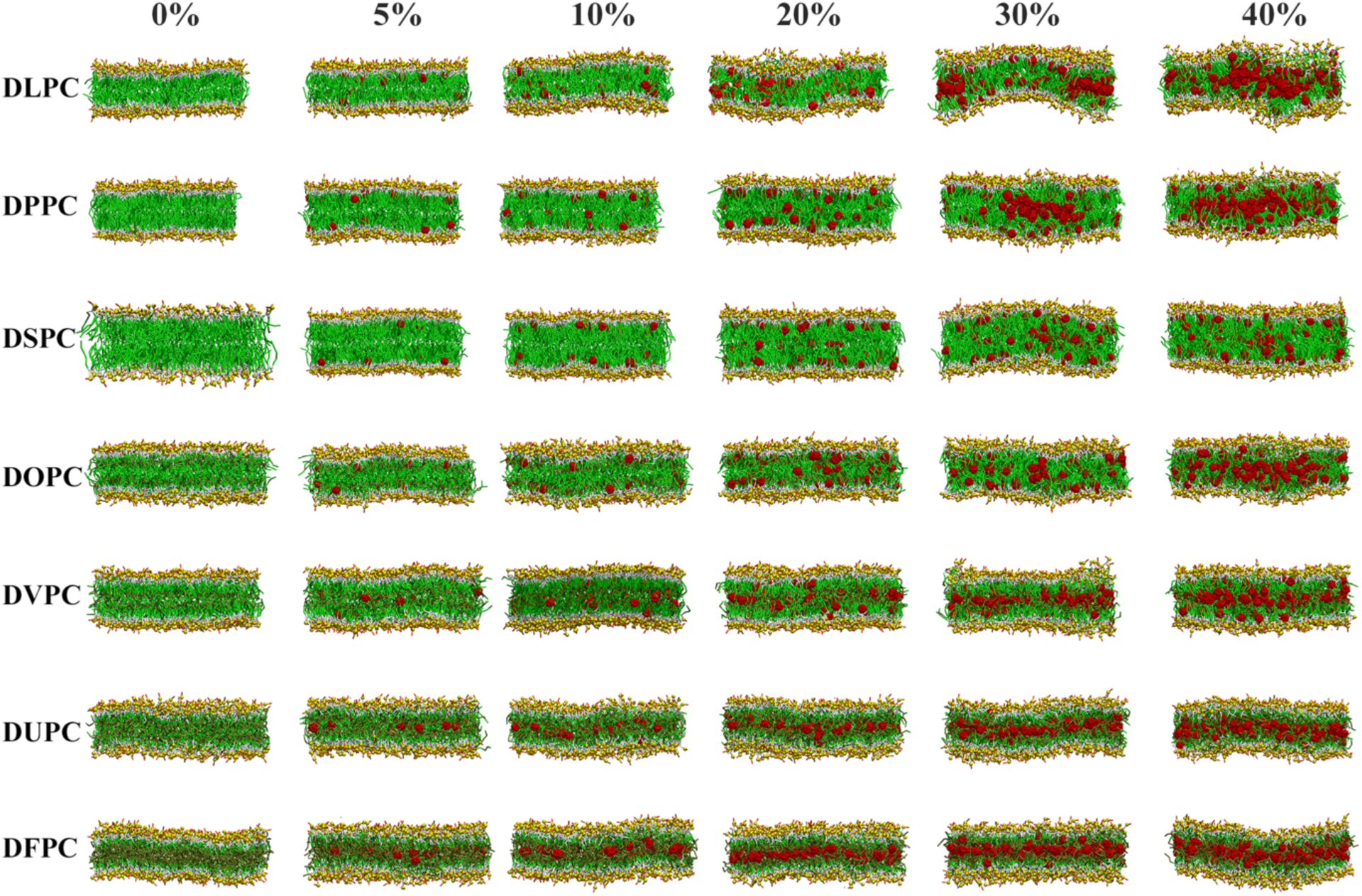
Snapshots illustrating the last frames of the systems in the *xz*-plane at different fullerene concentrations in the lipid bilayers (DLPC, DPPC, DSPC, DOPC, DVPC, DUPC and DFPC). Green corresponds to apolar beads in the lipid tails (normal: saturated carbons, dark: double bonds). Red, orange, yellow and white beads represent fullerene, choline, phosphate and glycerol groups, respectively.

**Figure S3.**
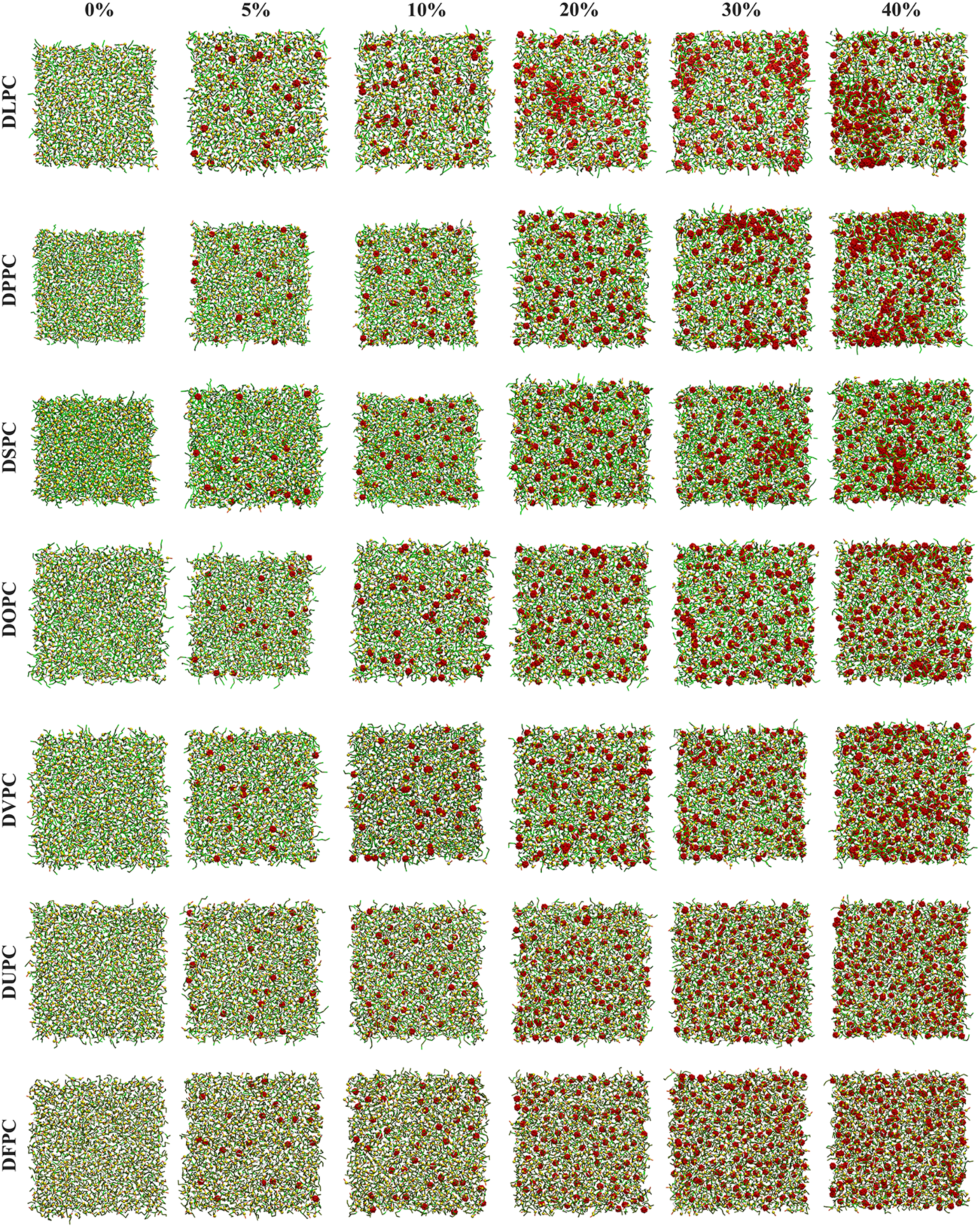
Snapshots illustrating the last frames of all systems in the *xy*-plane at different fullerene concentrations in the lipid bilayers (DLPC, DPPC, DSPC, DOPC, DVPC, DUPC and DFPC). Colors are as in Figure S1.

**Figure S4.**
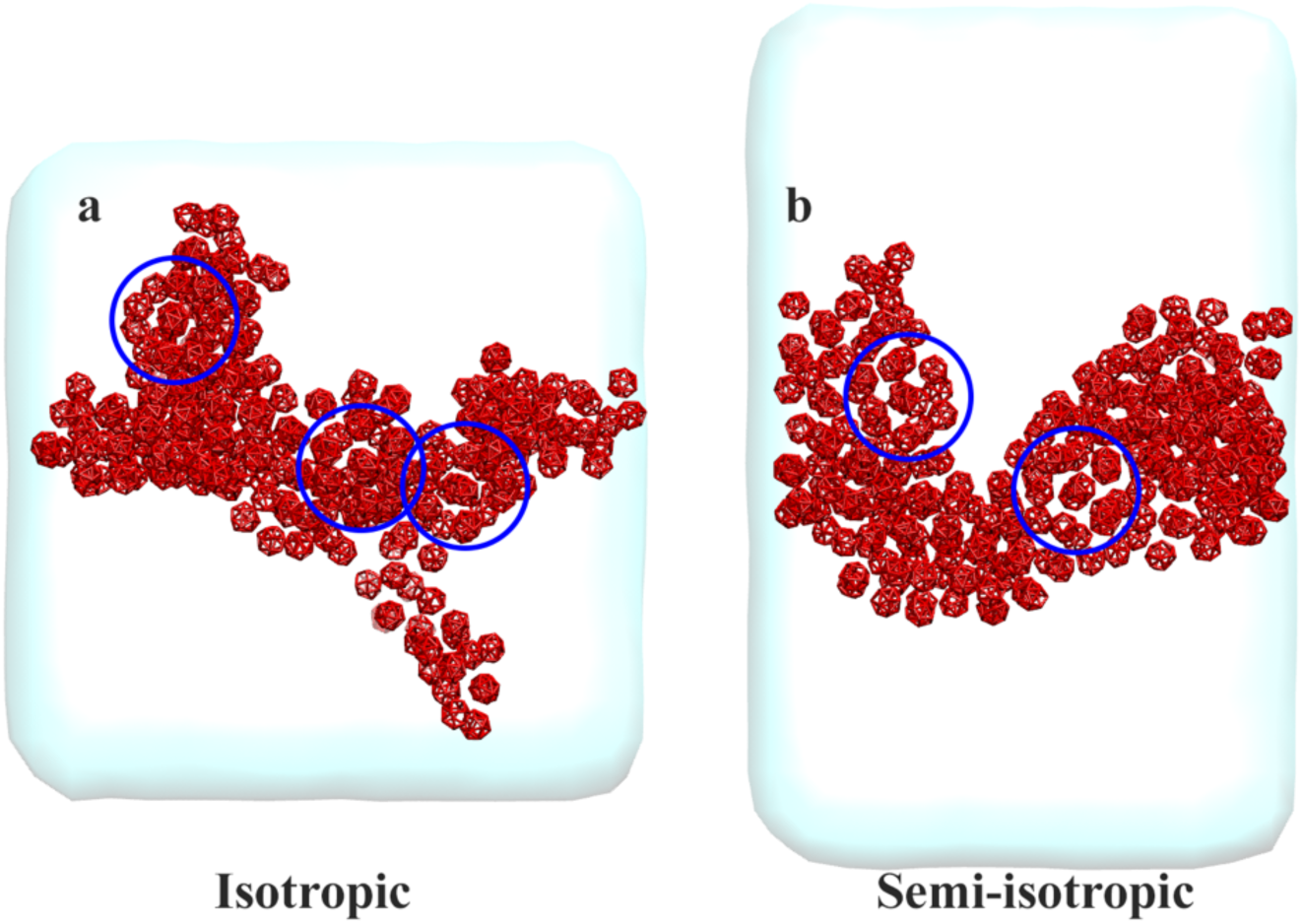
Final structures of C60 aggregates in water under two different pressure couplings (a) isotropic (b) semi-isotropic. Blue circles represent sphere-like clusters. Simulations were run for 5 µs.

**Table S2.** Coordination numbers of fullerene molecules were estimated by integrating the corresponding radial distribution functions (RDFs) up to 1.3 nm

|  | DLPC | DPPC | DSPC | DOPC | DVPC | DUPC | DFPC |
| --- | --- | --- | --- | --- | --- | --- | --- |
| 5% | - | - | - | - | - | - | - |
| 10% | 0.2 | 0.1 | 0.1 | 0.2 | 0.3 | 0.5 | 0.3 |
| 20% | 2.4 | 0.7 | 0.5 | 0.5 | 0.8 | 1.3 | 1.2 |
| 30% | 3.3 | 2.2 | 1.7 | 1.2 | 1.7 | 2 | 2.1 |
| 40% | 4.8 | 2.9 | 2.8 | 2.4 | 3 | 3 | 2.7 |

**Table S3.** Structural properties of the different membrane systems.

| Lipid Bilayer | Fullerene concentration | Area per lipid (nm<sup>2</sup>) | Volume per lipid (nm<sup>3</sup>) | Thickness (nm) |
| --- | --- | --- | --- | --- |
| DLPC | 0% | 0.593±0.001 | 1.026±0.001 | 3.460±0.001 |
|  | 5% | 0.615±0.001 | 1.062±0.001 | 3.455±0.001 |
|  | 10% | 0.672±0.001 | 1.099±0.001 | 3.443±0.001 |
|  | 20% | 0.675±0.002 | 1.179±0.001 | 3.507±0.003 |
|  | 30% | 0.698±0.003 | 1.250±0.001 | 3.582±0.008 |
|  | 40% | 0.690±0.004 | 1.303±0.003 | 3.780±0.020 |
| DPPC | 0% | 0.582±0.001 | 1.231±0.001 | 4.228±0.001 |
|  | 5% | 0.611±0.001 | 1.273±0.001 | 4.165±0.001 |
|  | 10% | 0.632±0.001 | 1.309±0.001 | 4.144±0.001 |
|  | 20% | 0.684±0.001 | 1.399±0.001 | 4.091±0.002 |
|  | 30% | 0.707±0.001 | 1.468±0.001 | 4.153±0.009 |
|  | 40% | 0.730±0.001 | 1.544±0.001 | 4.230±0.008 |
| DSPC | 0% | 0.595±0.001 | 1.452±0.001 | 4.877±0.001 |
|  | 5% | 0.610±0.001 | 1.485±0.001 | 4.870±0.001 |
|  | 10% | 0.629±0.001 | 1.521±0.001 | 4.837±0.001 |
|  | 20% | 0.679±0.001 | 1.612±0.001 | 4.769±0.004 |
|  | 30% | 0.709±0.001 | 1.687±0.001 | 4.760±0.004 |
|  | 40% | 0.732±0.001 | 1.763±0.001 | 4.824±0.005 |

**Figure S5.**
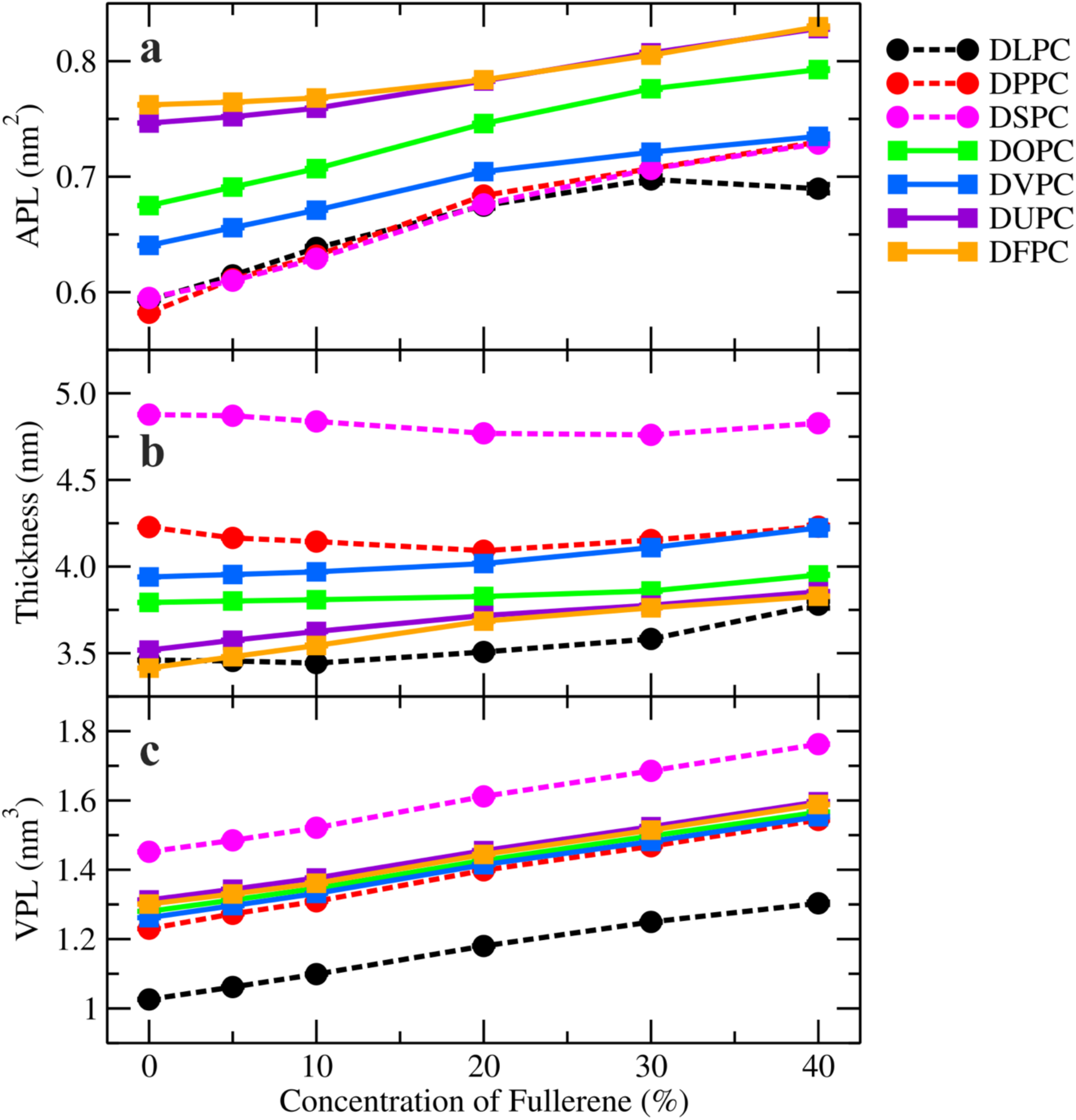
(a) Area per lipid (b) bilayer thickness (c) volume per lipid as a function of fullerene concentration. The dashed and solid lines correspond to saturated and unsaturated lipid bilayers, respectively. Error bars are smaller than the size of the symbols

**Figure S6.**
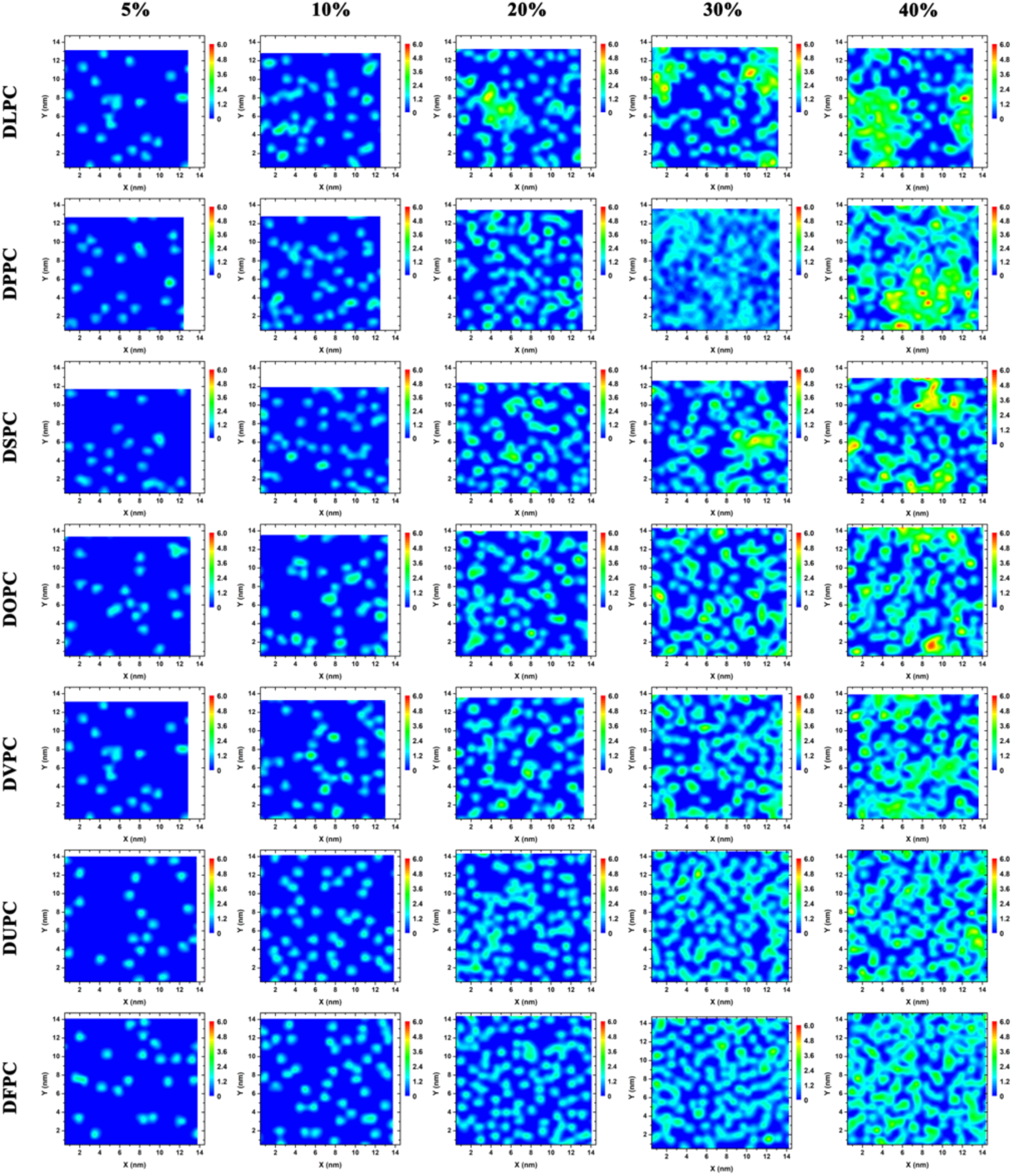
2D fullerene density maps at different fullerene concentrations.

**Figure S7.**
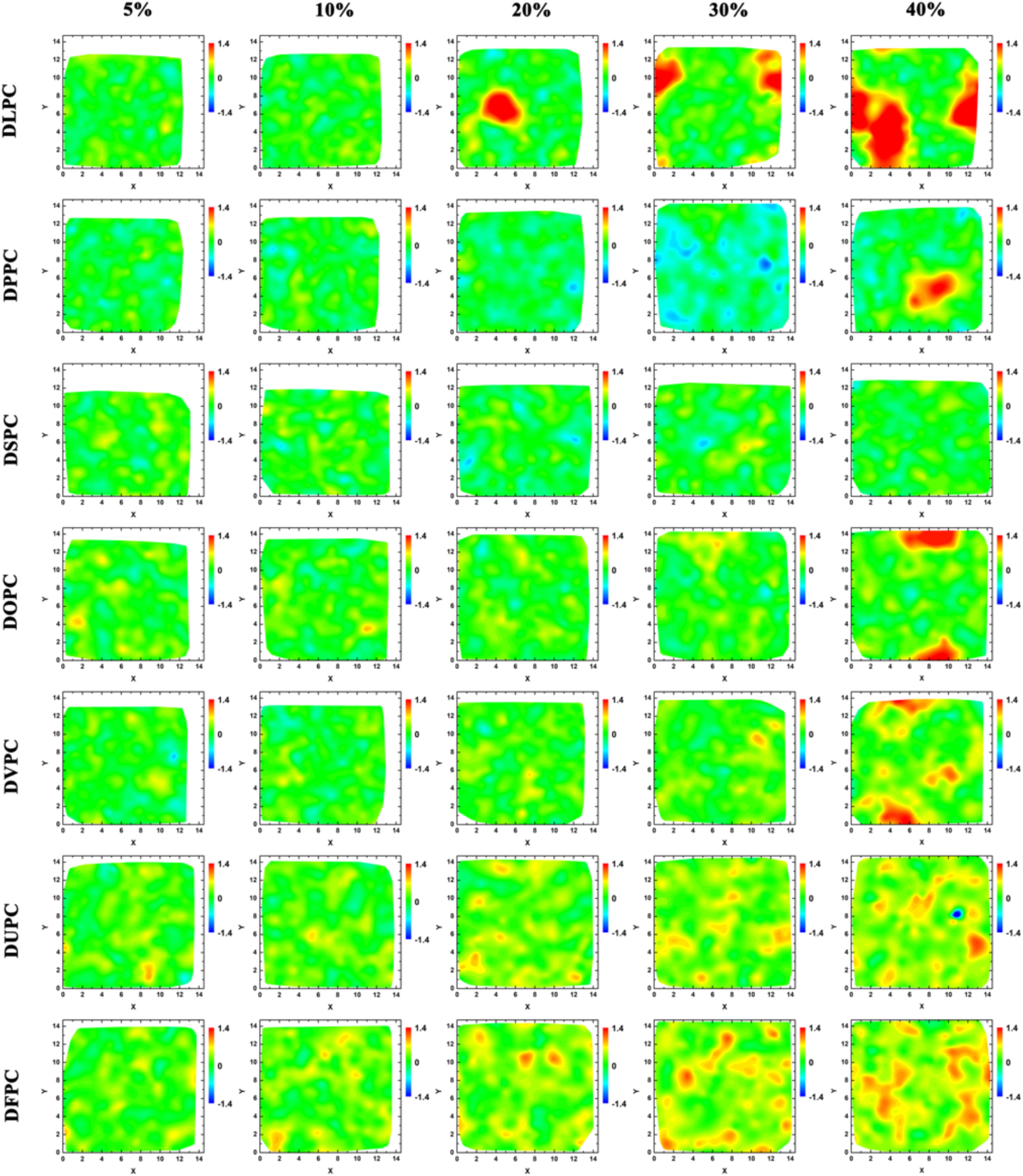
2D contour maps of bilayer thickness at different fullerene concentrations.

